# Predicting the physiological effects of multiple drugs using electronic health record

**DOI:** 10.1101/2024.06.30.601402

**Authors:** Junhyeok Jeon, Eujin Hong, Jong-Yeup Kim, Suehyun Lee, Hyun Uk Kim

## Abstract

Various computational models have been developed to understand the physiological effects of drug-drug interactions, which can contribute to more effective drug treatments. However, they mostly focus on interactions of only two drugs, and do not consider the patient information. To address this challenge, we use publicly available electronic health record (EHR), MIMIC-IV, to develop machine learning models that predict the physiological effects of two or more drugs. This study involves extensive preprocessing of laboratory measurement data, prescription data and patient data. The resulting machine learning models predict potential abnormalities across 20 selected measurement items (e.g., concentrations of metabolites and blood cells) in the form of a sentence. The models showed an average AUROC of 0.75, and age, specific active pharmaceutical ingredients, and gender appeared to be the most influential features. The model development process showcased in this study can be extended to other measurement items for a target EHR.

## INTRODUCTION

Taking multiple drugs simultaneously can lead to unexpected pharmacological effects that do not occur with individual drug use. These effects can maximize synergistic benefits when treating diseases, but also cause unexpected adverse drug events^1^. Consequently, understanding the effects of multiple drugs has become critical. In recent years, machine learning models have been developed to predict the effects of taking multiple drugs, often drug-drug interactions (DDIs), by processing various types of information^2,3,4^. DeepDDI^5^ and DeepDDI2^6^ use SMILES of two input drugs to predict their DDI effects by using deep learning. SSI-DDI^7^ employed molecular graphs to extract substructures of drug molecules, while DDIMDL^8^ and MDF-SA-DDI^9^ considered targets, pathways, and enzymes, as well as molecular features of drugs.

Despite advances in the DDI prediction, two main issues remain with these machine learning models. First, they mostly focus on interactions between only two drugs. However, in the real-world, patients often take more than two drugs simultaneously. For example, the average number of drugs taken by chronically ill patients recruited from 2019 through 2021 in Belgium was 5.7 with up to 20 drugs^10^. Similar observations were made for the Korean population, including the Korean outpatients^11^. Second, existing DDI-predicting machine learning models do not consider individual patient information. Drug responses can vary across different biological conditions. For example, age-related physiological changes^12^ as well as environmental and genetic factors^13^ can cause differences in drug responses across different ethnic and gender groups. Therefore, considering patient information is also important when predicting the physiological effects of multiple drugs.

In this study, we developed machine learning models to predict the physiological effects of multiple drugs by using three data within MIMIC-IV dataset: laboratory measurement data, prescription data, and patient data (Figure 1). MIMIC-IV dataset contains medical data from patients admitted to Beth Israel Deaconess Medical Center (BIDMC) between 2008 and 2019^14^. By using the MIMIC-IV dataset, this study aimed to develop machine learning models that predict the physiological effects of two or more drugs while considering patient information, thus addressing the two abovementioned problems of DDI-predicting models. Here, the physiological effects correspond to changes in measurement items (e.g., concentrations of metabolites and blood cells) selected from MIMIC-IV. The resulting machine learning models predicted potential abnormalities across 20 selected measurement items by using patient data (age, gender, and ethnicity) and prescription data as input (Figure 1). The output is generated in the form of a sentence. Influential input features were also analyzed where age appeared to be the most influential feature. Establishing the data preprocessing workflow was critical for developing the machine learning models. This study suggests how to utilize electronic health records (EHRs) to predict a patient’s response to multiple drugs, providing a model development process for predicting physiological effects that can be applied to other measurement items.

**Figure 1.**
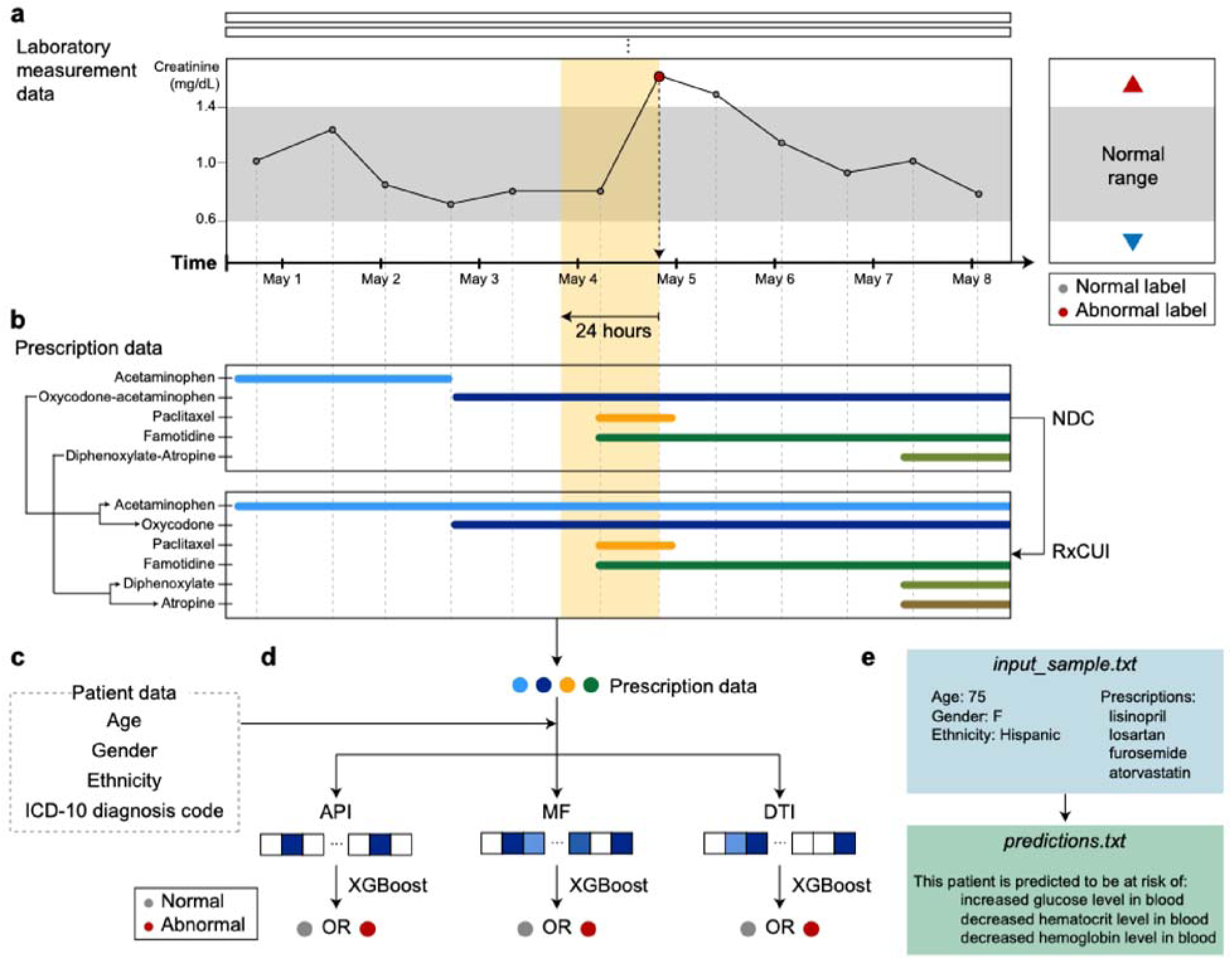
Overall scheme for the development of machine learning models to predict the physiological effects of multiple drugs. Laboratory measurement data (**a**), prescription data (**b**), and patient data (**c**) from MIMIC-IV were used to develop XGBoost models that classify the effects of multiple drugs on the measurement items (d). (**a**) Measurement instances of creatinine for a single *hadm_id* (i.e., a single patient) as an example. (**b**) Prescription data from the same patient in (**a**), which are aligned with the measurement instances across the timestamps. Here, all drugs prescribed within the preceding 24 hours of each measurement timestamp were considered (yellow box). Drug information presented using National Drug Codes (NDCs) were converted to RxNorm Concept Unique Identifiers (RxCUIs) of their corresponding active pharmaceutical ingredients (APIs) during preprocessing of the prescription data (**c**) Patient data considered in this study. (**d**) Three ways of considering the APIs when developing the machine learning models. Presence or absence of APIs, molecular fingerprints (MFs), and drug-target interactions (DTIs) were separately considered as input features to develop the models. The resulting XGBoost models predict the physiological effects of multiple drugs across the 20 selected measurement items. (**e**) An example of inputs and outputs of an API-based XGBoost model. The output is generated in the form of sentence.

## RESULTS

### Preprocessing of laboratory measurement data

Laboratory measurement data, mostly blood test data, was available in *labevents.csv.gz* from the MIMIC-IV dataset^14,15^. This laboratory measurement data was subjected to the preprocessing workflow, consisting of three data reduction steps (Figure 2a,b) and a labeling step (Figure 2c). The first two data reduction steps remove unclear measurement instances (i.e., unclear *charttime* values), and select measurement items (e.g., concentrations of creatinine and urea nitrogen, and counts of red blood cells in blood), based on the numbers of measurement instances and patients (Figures 2a,b). As a result of these two reduction steps, top 20 measurement items were selected as the target physiological effects of multiple drug administration (Figure 2b). The measurement items starting from the rank 21^st^ started showing substantially lower values for both the numbers of measurement instances and patients. Third, for each measurement item, patients who already had abnormal measurement data from the start of their hospitalization were not considered (Figure 2a). However, patients with at least one abnormal measurement result during their hospitalization were considered for each measurement item.

**Figure 2.**
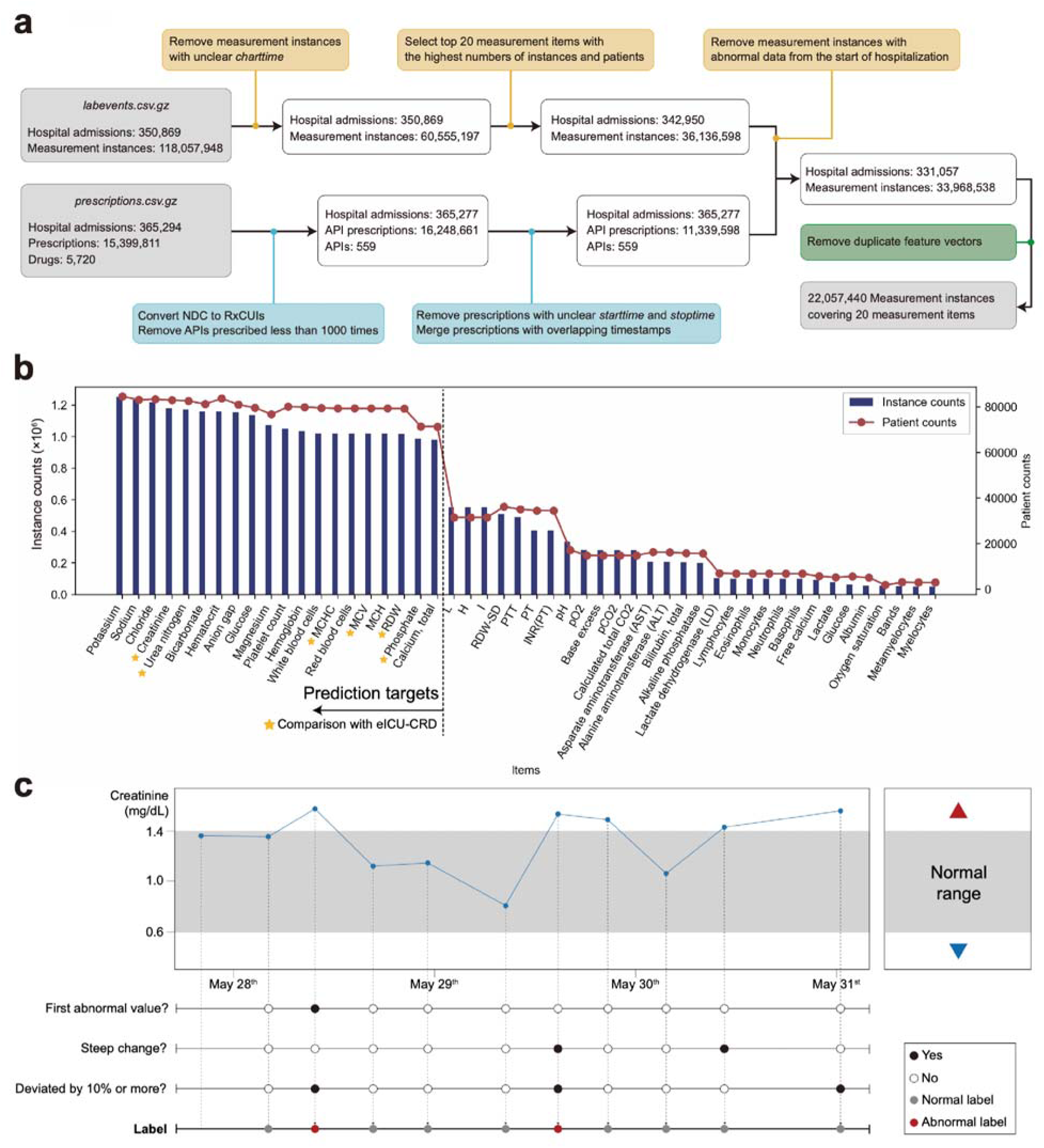
Preprocessing of the MIMIC-IV dataset. (**a**) Statistics of the laboratory measurement data (top) and the prescription data (bottom) from MIMIC-IV during the preprocessing. The preprocessed laboratory measurement data and prescription data were integrated into one dataset at the end, resulting in 22,057,440 measurement instances. (**b**) The numbers of measurement instances and patients across the measurement items available in MIMIC-IV. Measurement items were sorted in descending order, and the 20 measurement items were selected to develop machine learning models that predict the physiological effects of multiple drugs. Measurement items with star marks are the ones also covered by eICU-CRD dataset (Figure 5). (**c**) Criteria for labeling the effects of multiple drugs on the laboratory measurement data. A laboratory measurement instance was labeled as abnormal if any two of the three presented conditions were met. The last line “Label” presents the final label results (“Normal label” or “Abnormal label”) for each measurement instance.

The selected measurement instances were subsequently labeled (Figure 2a,c). A given measurement value was labeled as abnormal if at least any two of the following conditions were met: (1) the first abnormal value (*valuenum*) for a measurement item upon admission; (2) a steep increase or decrease (top 50% of all the change rates) for a measurement value in comparison with its previous value (*valuenum*); and (3) deviation from a normal range by 10% or more than 10% of the normal range. As a result of this preprocessing, from 118,057,948 measurement instances covering 350,869 hospital admissions (*hadm_id*) for the entire measurement items, a total of 36,136,598 measurement instances were finally selected for the 20 measurement items (Figure 2a). The selected data were used to develop machine learning models for each of the 20 measurement items. It is important to note that a single patient may have multiple hospital admissions, and the patients may have different conditions with each admission.

### Preprocessing of prescription data

Prescription data available in *prescriptions.csv.gz* from MIMIC-IV were also subjected to systematic preprocessing after many trials (Figure 2a). Drugs that correspond to *MAIN* and *ADDITIVE* types were considered from *prescriptions.csv.gz*; those belonging to *BASE* type (e.g., solvents or carriers commonly used for other drugs) were not considered. While preprocessing the prescription data, we decided to use RxNorm Concept Unique Identifiers (RxCUIs) for APIs rather than National Drug Codes (NDCs) for drug products. Different drug products are assigned with different NDCs despite having the same APIs; often, a drug product contains multiple APIs. This means that different drug products containing the same APIs can appear multiple times, leading to unnecessary computations without much contribution to the model performance. Indeed, XGBoost^16^ models developed by using RxCUI and NDC showed comparable performances, but the former took much shorter time for training. Also, using RxCUI makes it more convenient to handle drug information than NDC because one RxCUI indicates one API. Next, drugs prescribed less than 1000 times in *prescriptions.csv.gz* were further removed; this removal resulted in 559 APIs (or RxCUIs) from 1592 APIs (Figure 2a). Prescription data were also removed if their timestamps were not available, including those with unclear *starttime* or *stoptime* values. Finally, prescription data were merged if they had overlapping timestamp. It should be noted that dose value information was not considered because machine learning models did not benefit significantly from its inclusion.

With the preprocessed prescription data, API data were handled in three different ways as input features in this study. They were: a 559-dimensional vector presenting the presence or absence of APIs, a 1,024-dimensional vector for molecular fingerprints (MFs), and a 2,472-dimensional vector for drug-target interactions (DTIs). The 2,472-dimensional vector specifically contains information on target proteins and interaction modes of an API. For example, target proteins of furosemide are solute carrier family 12 member 1 inhibitor and G protein-coupled receptor 35 agonist. Examples of the interaction modes considered in the 2,472-dimensional DTI vector include inhibitor, activator, substrate, product, ligand, inducer, antibody, cleavage, metabolizer, cofactor, binder, agonist, antagonist, potentiator, and blocker. Consequently, three machine learning models were developed for each of the 20 measurement items, each using one of the three API features (Figure 3). For convenience, the three types of models were referred to as “API-based models”, “MF-based models”, and “DTI-based models”.

**Figure 3.**
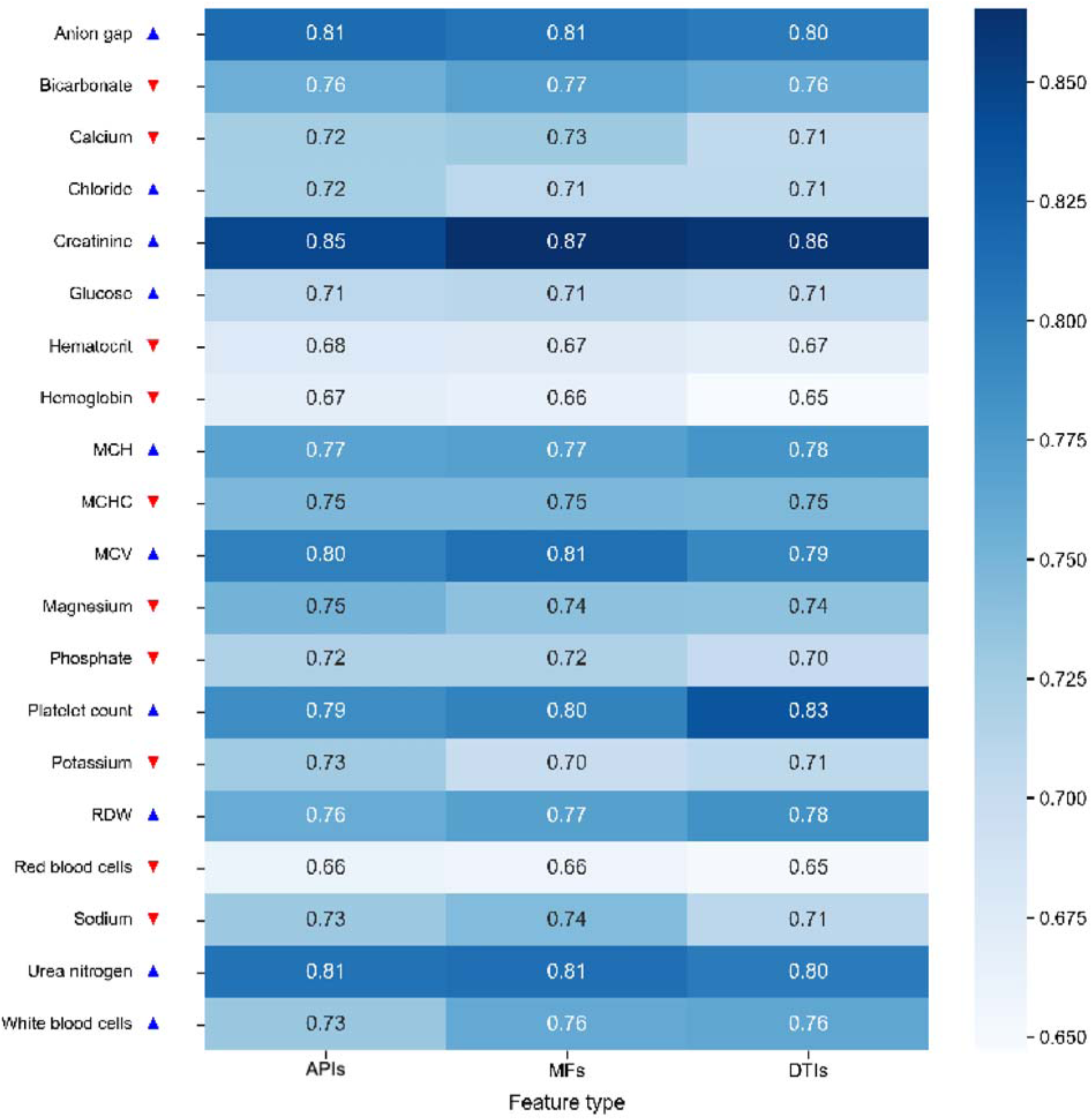
AUROC of the machine learning models developed for the 20 measurement items. Three different types of machine learning models were developed for each measurement item by using: information on presence or absence of active pharmaceutical ingredients (APIs); molecular fingerprints (MFs) of APIs; and drug-target interactions (DTIs) of APIs.

### Preparation of datasets to develop machine learning models

To develop the three types of machine learning models for each measurement item, the labeled measurement data (Figure 2c) were integrated with the preprocessed prescription data and the patient data (Figures 1 and 2a). For each measurement label (i.e., normal or abnormal) at a specific timestamp, all APIs prescribed within the preceding 24 hours of that timestamp were considered (Figure 1). The patient data includes gender, age, and ethnicity (Figure 1). After this step, a dataset containing 35,954,346 instances across the 20 measurement items has been prepared for 333,599 hospital admissions (Figure 2a).

The final step for the dataset preparation was to remove duplicate feature vectors that contain information on the drug and patient data (Figure 2a). When patients are prescribed medication, they typically take the same drug for several days, generating the same feature vectors within and across patients. To prevent redundancy, only one instance was retained, while removing all the same redundant instances. The selected data were labeled as abnormal if there was at least one abnormal measurement instance across the duplicate data; otherwise, the selected data was labeled as normal. It should be noted that the duplicate feature vectors may have different measurement labels. As a result, a dataset of 22,057,440 instances was prepared for the 20 measurement items (Figure 2a). This dataset was subsequently used to prepare independent training, validation and test datasets, at the ratio of 7:1:2, for each of the 20 measurement items, each covering 305,270 hospital admissions and 1,470,496 measurement instances on average.

### Development of machine learning models for predicting the effects of multiple drugs

Using the prepared datasets, machine learning models were developed for each measurement item. Model input consists of patient data and prescription data (Figure 1e). Patient data include aga, gender, and ethnicity. Prescription data indicate prescribed drugs, which are represented using information on the presence or absence of APIs, MFs, or DTIs. Therefore, one has to choose which type of model to run among the API-based model, MF-based model, and DTI-based model, depending on the input features of APIs. The prediction outcome is a sentence indicating potential abnormalities across the 20 selected measurement items (Figure 1e).

As a result, the three types of models showed similar classification performance by using the test datasets for the 20 selected measurement items. The average AUROC was 0.75 for the API-based models, 0.75 for the MF-based models, and 0.74 for the DTI-based models (Figure 3). Overall, classification for the creatinine measurement showed the best performance with an AUROC range of 0.85 to 0.87, followed by the anion gap and urea nitrogen measurements, both having an AUROC range of 0.80 to 0.81 (Figure 3). Meanwhile, red blood cells and hemoglobin were the least predictable items, both showing an AUROC range of 0.65 to 0.67.

### Analysis of the effects of input features on prediction outcomes

The importance of input model features was analyzed using SHapley Additive exPlanations (SHAP)^17^ (Figure 4). SHAP value for each feature indicates its relative impact on a model output. A more positive SHAP value indicates a more positive impact on a model output, and a negative SHAP values refers to the opposite meaning (i.e., more negative impact). In our models, higher SHAP values correlate with a higher probability of having abnormality for a given measurement item. The top 20 features with the highest mean absolute SHAP values were extracted for each model (Figure 4). As a result, the most important feature was age in 16 out of the 60 machine learning models developed for the 20 measurement items. Furosemide appeared to be the second most influential feature after age in the API-based model developed for urea nitrogen (Figure 4a). Furosemide is an API used to treat edema and has a known adverse event of hyperuricemia^18^. The association between urea nitrogen and furosemide, with a focus on hyperuricemia, could be further studied. This way of analysis can be extended to other pairs of measurement items and prescribed drugs.

**Figure 4.**
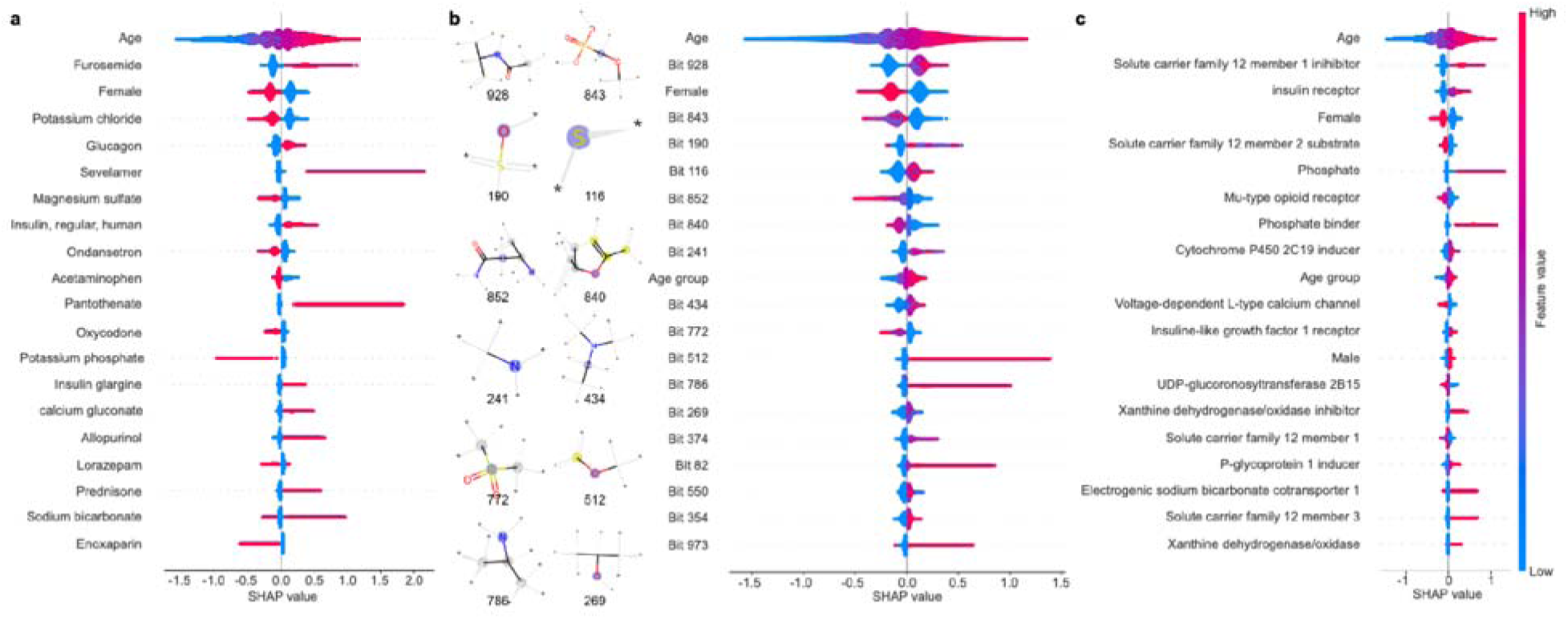
SHAP values of top 20 input features for the machine learning predictions for urea nitrogen. Top 20 input features are presented based on their highest mean absolute SHAP value for the API-based model (**a**), MF-based model (**b**), and DTI-based model (**c**). SHAP value of each feature was calculated for each measurement instance. Each data point represents a single prediction, and its color indicates positive or negative impact of the feature on the model’s prediction. SHAP values of each feature were independently scaled.

Notably, the predictions made by the three types of models, using APIs, MFs, and DTIs, were consistent with one another in general (Figure 4). For example, solute carrier family 12 member 1 (from the DTI-based model; Figure 4c) is a target protein for furosemide and potassium chloride (from the API-based model; Figure 4a), while acetaminophen and lorazepam (from the API-based model; Figure 4a) target UDP-glucuronosyltransferase 2B15 (from the DTI-based model; Figure 4c). Likewise, furosemide and insulin glargine (from the API-based model; Figure 4a) share a molecular substructure “Bit 928” (from the MF-based model; Figure 4b), and glucagon, pantothenate, insulin glargine, and calcium gluconate (from the API-based model; Figure 4a) share “Bit 434” (from the MF-based model; Figure 4b). These examples demonstrate the value of using different API features through independent machine learning models, thereby facilitating the analysis of prediction outcomes.

### External validation of the machine learning models using eICU-CRD dataset

To ensure the robustness of the models, they were also validated using eICU-CRD dataset^19,20^ (Figure 5). The models developed for the following six selected measurement items were subjected to the external validation: bicarbonate, chloride, mean corpuscular hemoglobin concentration (MCHC), mean corpuscular volume (MCV), phosphate, red blood cell distribution width (RDW), and urea nitrogen in blood (Figure 5a). Normal ranges for these six measurement items appeared to be the same across all the patients in the MIMIC-IV dataset. These normal range values were used to label the measurement data from eICU-CRD; eICU-CRD data have measurement data (i.e., measurement instances), but do not have a defined normal range for each measurement instance. It should be noted that the other 14 measurement items considered in this study (Figure 3) have different normal ranges in the MIMIC-IV dataset, depending on patients. Laboratory measurement data and prescription data from the eICU-CRD dataset were also subjected to preprocessing (Figure 5b). As a result of comparison with the preprocessed eICU-CRD data, all the models showed classification performances with a median AUROC of 0.66 across the six measurement items (Figure 5c). This external validation suggests that the models still retain their predictive power despite being even when applied to data from a different source.

**Figure 5.**
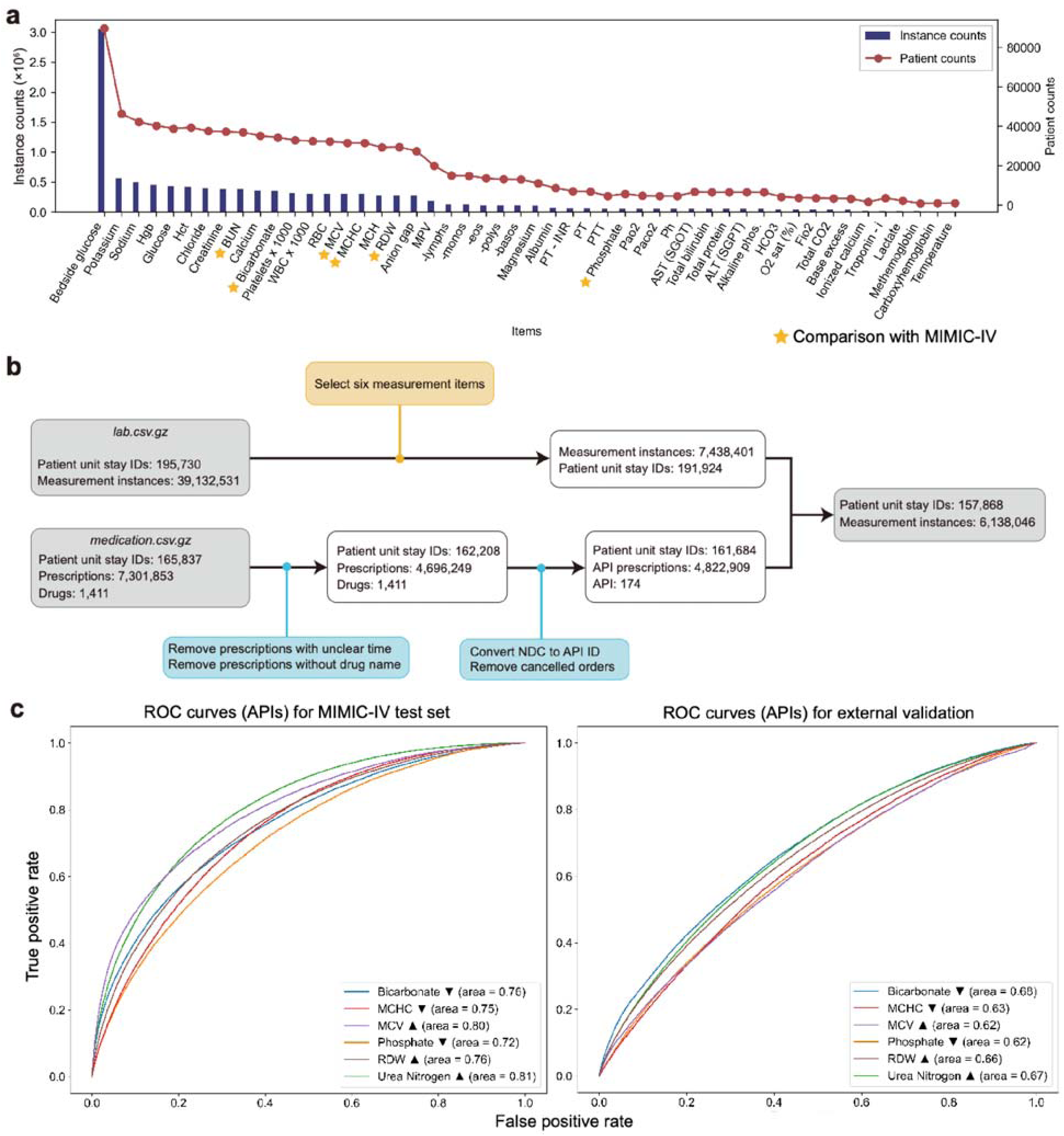
External validation of the machine learning models using the preprocessed eICU-CRD dataset. (**a**) The numbers of measurement instances and patients across the measurement items in eICU-CRD. Measurement instances for the starred measurement items were used to validate the machine learning models developed using MIMIC-IV. (**b**) Statistics of the laboratory measurement data (top) and the prescription data (bottom) from eICU-CRD during the preprocessing. The preprocessed laboratory measurement data and prescription data were integrated into one dataset at the end, resulting in 6,138,046 measurement instances. (**c**) Receiver operating characteristic (ROC) curves for the machine learning models developed using active pharmaceutical ingredients (APIs) for the six selected measurement items.

## DISCUSSION

In this study, machine learning models were developed to predict patients’ physiological responses to multiple drugs. When developing machine learning models, this study attempted to address the two challenges: consideration of three or more drugs, and individual patient data (Figure 1). These challenges could be addressed by using the EHR dataset, MIMIC-IV. However, the MIMIC-IV dataset has a highly complex structure, and required systematic preprocessing to prepare training datasets for machine learning models (Figure 2). As a result, the developed models successfully predicted outcomes across the 20 measurement items (Figures 3 and 5). Importantly, the API-based, MF-based, and DTI-based models were used to generate prediction outcomes, which were analyzed together to identify important input features. For example, age, specific active pharmaceutical ingredients, and gender appeared to be the most influential features when predicting the effects of multiple drugs on the urea nitrogen measurement item.

This study has also revealed limitations and suggestions for future studies. First, the broad scope of measurement items required comprehensive preprocessing of the EHR datasets. During the preprocessing, the assignment of abnormal labels to measurement instances was not always medically clear, and has room for further exploration. Second, the diversity of the patient data is limited. Both the MIMIC-IV and eICU-CRD datasets used for training and validation were collected from U.S. hospitals. Furthermore, the datasets come from intensive care unit (ICU) patients. Consequently, the prepared measurement data has a higher proportion of abnormal labels than might be typical in non-ICU settings. This is particularly true for certain measurements that are commonly abnormal in ICU environments, such as high glucose, low hematocrit, low hemoglobin, and low red blood cells all considered in this study (Figure 3).

Future studies for predicting the physiological effects of multiple drugs should carefully consider these limitations. If properly addressed, machine learning models could be used to also detect adverse drug reaction signals as reported previously^21,22,23^. Also, the preprocessing workflow of EHR datasets can be applied to other measurement items to capture more diverse physiological effects of multiple drugs.

## METHODS

### Ethical approval and consent

The institutional review board of KAIST approved the waiver of consent for individual patients because this study was retrospective and all the data used in this study were de-identified (IRB No. IRB-22-230). For MIMIC-IV, the collection of patient information and creation of the research resource was reviewed by the Institutional Review Board at the Beth Israel Deaconess Medical Center, who granted a waiver of informed consent and approved the data sharing initiative. For eICU-CRD, the study is exempt from institutional review board approval due to the retrospective design, lack of direct patient intervention, and the security schema, for which the re-identification risk was certified as meeting safe harbor standards by an independent privacy expert (Privacert, Cambridge, MA) (Health Insurance Portability and Accountability Act Certification no. 1031219-2).

### Collection and preprocessing of the MIMIC-IV and eICU-CRD datasets

Within MIMIC-IV v2.1, following six datasets were initially used in this study^14,15^: *labevents.csv.gz* and *d_labitems.csv.gz* for laboratory measurement data of ICU patients; *prescriptions.csv.gz* for prescription data involving the use of NDC; *patients.csv.gz* and *admissions.csv.gz* for patient data; and *diagnoses_icd.csv.gz* for diagnosed diseases using ICD-10 diagnosis code. *lab.csv.gz* and *medication.csv.gz* were used for measurement data and prescription data, respectively, from the eICU-CRD dataset^19,20^. Each of these tabular datasets underwent different processing procedures (Figures 2a and 5b). ICD-10 diagnosis code in *diagnoses_icd.csv.gz* was initially considered, but later no longer considered. Additional resources were used to process the prescription data, including RxNorm API 3.1.254 and RxNav^24^ 2.9.119 and DrugBank^25^ 5.1.8, as well as Python packages such as RDKit 2023.03.1 (http://www.rdkit.org). RxNorm was used to convert NDC of a drug into its APIs (Figure 1). DTI information of APIs was obtained from DrugBank. MFs of APIs were generated using RDKit.

### Development of machine learning models

All the data processing and modeling were conducted on Python 3.11.3 using the following standard libraries: pandas^26^ 1.5.3, numpy^27^ 1.24.3, scipy^28^ 1.10.1, scikit-learn^29^ 1.2.2, xgboost^16^ 1.7.3, pytorch^30^ 1.7.1, pytorch-tabnet^31^ 4.0, matplotlib^32^ 3.7.1, seaborn^33^ 0.11.2, and shap^17^ 0.41.0.

To optimize the performance of the XGBoost models, the Bayesian optimization approach was employed using Optuna^34^. Following hyperparameter were examined for the designated ranges in the Bayesian optimization: learning_rate, 10^-4^ to 10^-1^, max_depth, 3 to 10, min_child_weight, 10^-3^ to 10^2^), subsample, 0.1 to 1, colsample_bytree, 0.1 to 1, reg_alpha, 10^-6^ to 10^2^, reg_lambda, 10^-6^ to 10^2^, and n_estimators, 50 to 300. During the Bayesian optimization process, an early stopping criterion was set to 20 rounds. The number of trials conducted for Bayesian optimization was set to 30. After the Bayesian optimization, a XGBoost model with the highest AUC was selected for further studies.

## DATA AVAILABILITY

The MIMIC-IV and eICU-CRD datasets used in this study can be accessed at https://physionet.org/content/mimiciv/2.1/ and https://physionet.org/content/eicu-crd/2.0/, respectively.

## CODE AVAILABILITY

Code for the 60 machine learning models developed for the 20 selected measurement items is available at https://github.com/kaist-sbml/multi-drug-response.

## Acknowledgements

This research was supported by: KAIST Key Research Institute (Interdisciplinary Research Group) Project; an internal fund/grant of Electronics and Telecommunications Research Institute (ETRI) [22RB1100, Exploratory and Strategic Research of ETRI-KAIST ICT Future Technology]; and Ministry of Science and ICT through National Research Foundation of Korea (RS-2023-00262527).

## Author information

H.U.K. conceived the project. J.J. and E.H. conducted all the computational analyses. All the authors analyzed the data. J.J., E.H. and H.U.K. wrote the manuscript.

